# Scholar Metrics Scraper (SMS): automated retrieval of citation and author data

**DOI:** 10.1101/2021.12.23.473883

**Authors:** Nicole A. Cheung, Dean Giustini, Jeffrey LeDue, Tim H. Murphy

**Affiliations:** University of British Columbia, Department of Psychiatry, 2211 Wesbrook Mall, Vancouver, V6T 1Z3, British Columbia, Canada; Djavad Mowafaghian Centre for Brain Health, University of British Columbia, Vancouver, British Columbia, Canada; University of British Columbia Biomedical Branch Library, Vancouver, British Columbia, Canada

**Keywords:** Google Scholar, Python, Bibliometrics, Citation Metrics, Research Impact, Automation

## Abstract

Academic departments, research clusters and evaluators analyze author and citation data to measure research impact and to support strategic planning. We created a tool, Scholar Metrics Scraper (SMS), to automate the retrieval of this bibliometric data for our research team. The project contains Jupyter notebooks (publicly-shared here) that take a list of researchers as an input to export a CSV file of citation metrics from Google Scholar and figures to visualize the group’s impact. SMS is a scalable, open and publicly-accessible solution for automating the retrieval of citation data over time for a group of researchers.

## 1. Motivation and significance

The most widely-accepted, objective metrics used to examine productivity and influence in science are linked to a researcher’s publications, and the number of times they are cited by others (Schreiber & Giustini, 2019). For example, the h-index combines two key metrics, publication and citation counts, into a single number that can be used to evaluate a researcher’s impact (Hirsch, 2005). Patterns comprising analysis of citation metrics support a range of scholarly activities from identifying collaborative performance and research trends to strategic planning. In fact, tenure and promotion committees, granting agencies and administrators spend considerable time evaluating researchers and the impact of their publications (De Silva & Vance, 2017).

Several indexes are used to perform bibliometric analysis and to extract accurate citation data over time. PubMed, produced by the US National Library of Medicine, is a key bibliographic database in biomedicine but it is not designed to extract citation data. For that purpose, tools such as Google Scholar (GS), Scopus and the Web of Science (WoS) are used and offer important ways to gather this information. However, Scopus and WoS require a subscription and limit the ways information can be extracted. For its part, GS holds a unique place due to its size and by providing a freely-accessible interface to acquire bibliometric data (Martín-Martín et al., 2020).

Third party tools such as Bibliometrix and Publish or Perish can retrieve citation data from several data sources (Aria & Cuccurullo, 2017; Harzing, 2007), but are not designed to retrieve data for research groups. Elsevier’s SciVal and Clarivate’s InCites are both proprietary research assessment tools used to access author metrics from the Scopus and WoS databases, respectively (Bornmann & Leydesdorff, 2013; Clarivate, n.d.; Dresbeck, 2015), but require subscription access and considerable skill to extract information efficiently. For programmers, Python packages such as pybliometrics and scholarly provide built-in methods to retrieve citation data from Scopus and GS (Cholewiak et al., 2021; Rose & Kitchin, 2019). Although these tools are becoming more widely-used in scientific departments, specific and efficient methods to locate bibliometric data for large groups of authors are needed.

Here, we present Scholar Metrics Scraper (SMS), a novel tool that uses the scholarly package to automate data extraction from GS for groups of researchers. In the form of Jupyter Notebooks (JN), it comes with step-by-step instructions for modification and interaction in a minimum number of steps without needing programming experience. SMS users can upload lists of authors and modify the JNs to retrieve a range of data types which are organized into a table. This table is used to create figures to track a research group’s collective scholarly output and to assess their impact and collaboration. To illustrate our SMS, we provide use cases within the Dynamic Brain Circuits in Health and Disease Research excellence cluster at the University of British Columbia.

## 2. Software Description

Definition of terms
|  |  |
| --- | --- |
| Scholar Metrics Scraper (SMS) | Our project contains the notebooks ScholarScraper, ScholarCollabs, and GroupedCollabs. It is available on GitHub at https://github.com/ubcbraincircuits/scholar_metrics_scraper. |
| ScholarScraper notebook | Contained in SMS as ScholarScraper.ipynb, it takes a list of authors as input, retrieves data from Google Scholar for each, and puts them into a table in an output CSV file. It also creates a citations/year bar chart and a collaboration heatmap as output. |
| ScholarCollabs notebook | Contained in SMS as ScholarCollabs.ipynb it takes the data table created by the ScholarScraper notebook and creates a collaboration chord diagram to visualize links between coauthors. |
| GroupedCollabs notebook | Contained in SMS as GroupedCollabs.ipynb it works similarly to ScholarCollabs, but groups authors into subgroups to create a grouped collaboration chord diagram. |
| Google Scholar (GS)<br>http://scholar.google.com | The web search engine is used to search for scholarly literature and provides access to citation and author metrics. Open and publicly-accessible, GS is the largest scholarly search engine on the web providing access to more than 300 million references. |
| scholarly | A Python package that provides built-in methods to retrieve author data from GS for a given researcher (Cholewiak et al., 2021) |

### 2.1 Software architecture

Our SMS project is available on GitHub at https://github.com/ubcbraincircuits/scholar_metrics_scraper, and comes with the ScholarScraper notebook (which retrieves the data from GS and creates a citations/year barchart and collaboration heatmaps) as well as the ScholarCollabs and GroupedCollabs notebooks (which create collaboration chord diagrams). Users are instructed to open Syzygy, which provides free online access to Jupyter (Lamoureux, 2020), and log in with their institution or Gmail account. A few steps are required to set up the project in Syzygy (see Fig 1 steps 1-3). Users can follow instructions to clone the project into their Jupyter directory (step 1) and install scholarly (step 2). Users create a CSV file with author names or GS identifiers (IDs) in a single column (see Appendix A) and upload the file to their Jupyter directory (step 3). A GS ID is alphanumeric and embedded within the GS profile’s uniform resource locator (URL). For example, where an author’s profile URL (Dr. Tim H. Murphy) is “https://scholar.google.ca/citations?user=qJjM8hkAAAAJ&hl=en“, the corresponding GS ID is qJjM8hkAAAAJ.

**Fig. 1.**
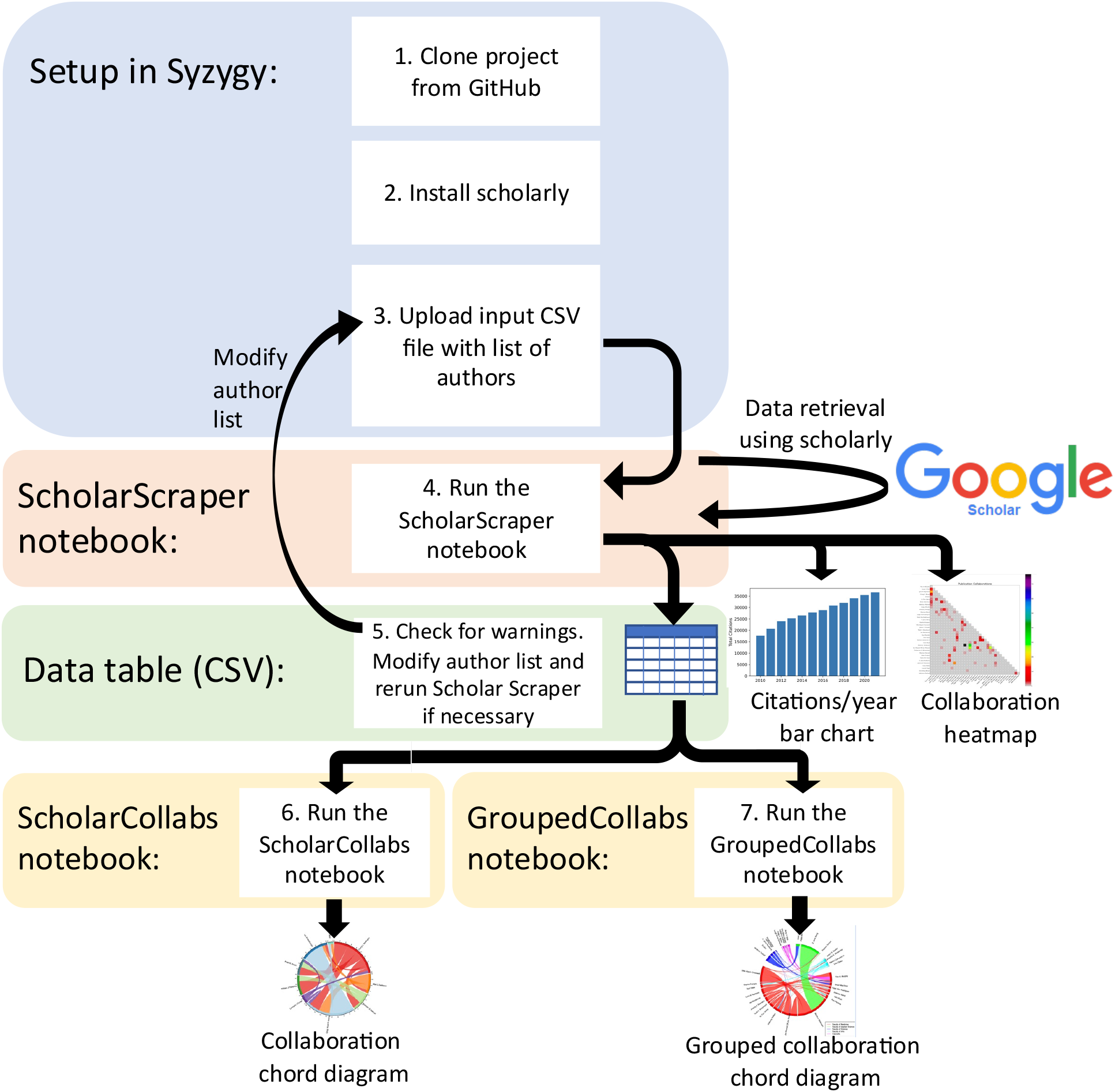
Scholar Metrics Scraper workflow diagram including the architecture and major steps.

### 2.2 Software functionalities

The ScholarScraper notebook contains Python code cells with step-by-step instructions to load in the author list, retrieve data from GS, and produce a table and two figures (see Fig 1 step 4). Users make minor modifications to the code in their author list and provide a list of institution names with which the authors are collectively affiliated (the software finds matches between author and institution lists). The ScholarScraper notebook iterates over the list of authors to retrieve the data from GS and output a data table as a CSV file. The output table will contain data such as publication and citation counts, h-index, publication titles and coauthors (see Appendix B). The notebook is flexible and can be modified to retrieve specific types of data or extract them for specific analyses.

Scholarly can use an author’s name to retrieve the closest match; so, a different author with a similar name may be found. To inform users when (or if) the retrieval is unsuccessful, the output table includes a warning column (see Appendix B). The ScholarScraper notebook checks each author’s affiliation (retrieved from GS) against the affiliations list as modified by the user. When none of the strings match the author’s affiliation, a message that the affiliation did not match will appear in this column. Users should check that names in the author list match their GS profiles and output data. As necessary, users can modify author names in the CSV file to ensure the correct data is collected (see Fig 1 step 5). Alternatively, the notebook can take less ambiguous GS profile IDs directly as an input instead of names.

Scholarly will not retrieve data for researchers unless they have already created a GS profile. Scholars can set up their own profiles in GS by manually adding their biographical information, expertise and keywords, and curating their publications (Thoma & Chan, 2019). If scholarly does not find an existing profile under the author’s name, it will output an error resulting in a blank row in the output table, and a warning will be given that no profile was found.

The ScholarScraper notebook takes each author’s citations each year over time and totals them to create a bar chart of total citations per year for the group (Fig. 2). When two authors in the group have a common publication as collaborators, citations for this publication will be double-counted. Therefore, yearly citation totals represented by the bar chart may overcount the group’s citations.

The ScholarCollabs notebook will also create a heatmap of the count of collaborations between coauthors (Fig 3). The heatmap is created using the data shown in Appendix C.

The project contains a ScholarCollabs notebook and a GroupedCollabs notebook which creates collaboration diagrams with step-by-step instructions and R code cells (see Fig 1, steps 6-7). After running the ScholarScraper notebook and creating an output data table as CSV, users can open the ScholarCollabs notebook, and follow instructions to modify the file name objects, title, and indicate if they want a weighted or unweighted diagram. The ScholarCollabs notebook takes the names of the authors, their coauthors, and the number of publications they share with each coauthor and creates a chord diagram to visualize co-authorships as links between authors (Fig. 4). The diagram is saved in the Jupyter directory as a PDF which can be downloaded locally. The GroupedCollabs notebook also creates collaboration diagrams, but produces a diagram with authors grouped into subgroups (e.g., faculty, research area, etc.) (Fig. 5). It works similarly to ScholarCollabs but takes an additional CSV file as input which contains the group for each author.

## 3. Illustrative Example

**Fig. 2.**
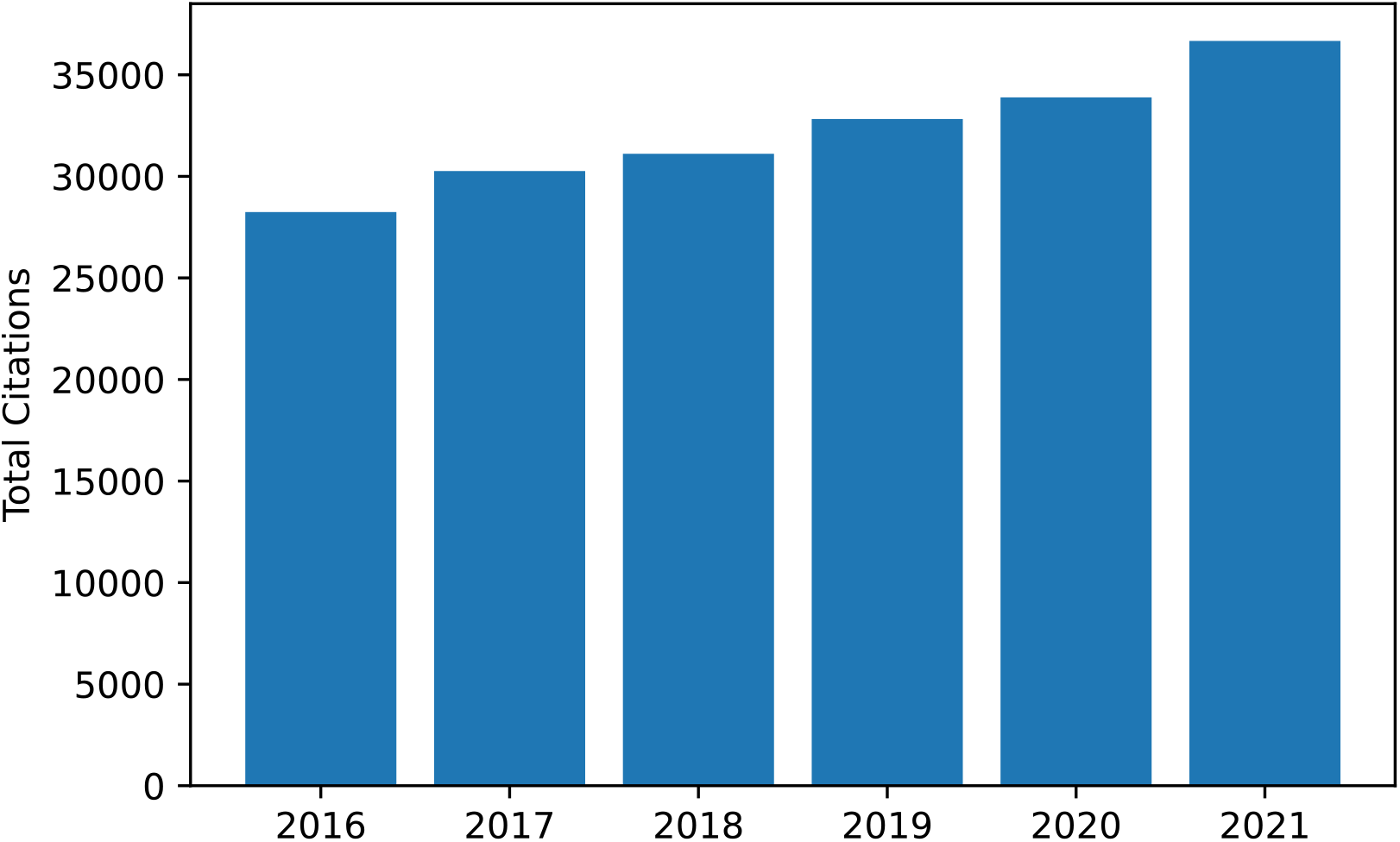
Total citations per year from 2016 to 2021. This includes citations for publications by all authors in the author list for the University of British Columbia Dynamic Brain Circuits in Health and Disease research Excellence cluster. This diagram is created as output by the ScholarScraper notebook, and the range of years can be easily modified. The graph reflects the total citations for the cluster and thus may contain duplicates within collaborative projects.

**Fig. 3.**
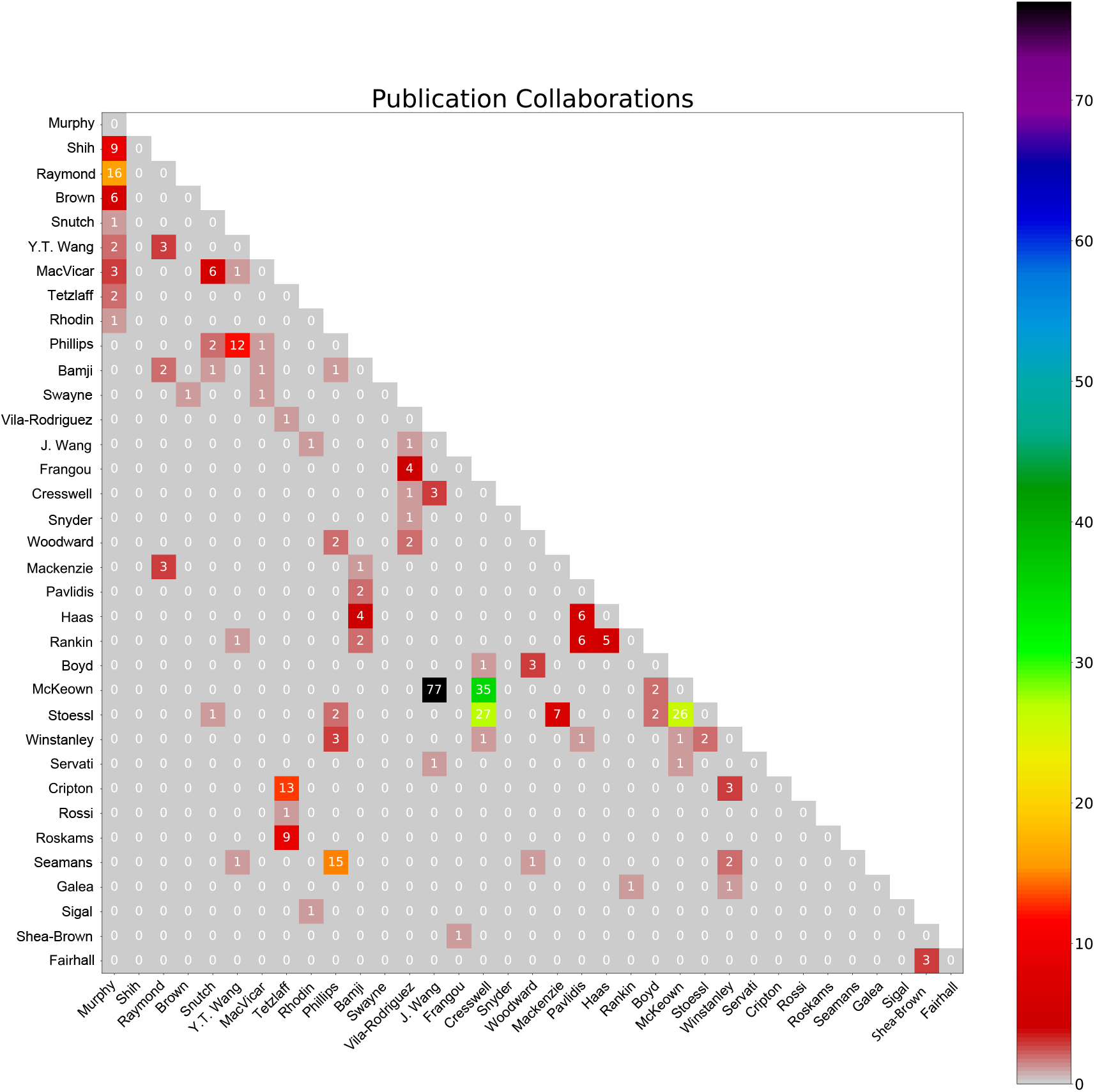
Count of collaborations heatmap. The numbers represent the count of publication collaborations between coauthors.

**Fig. 4.**
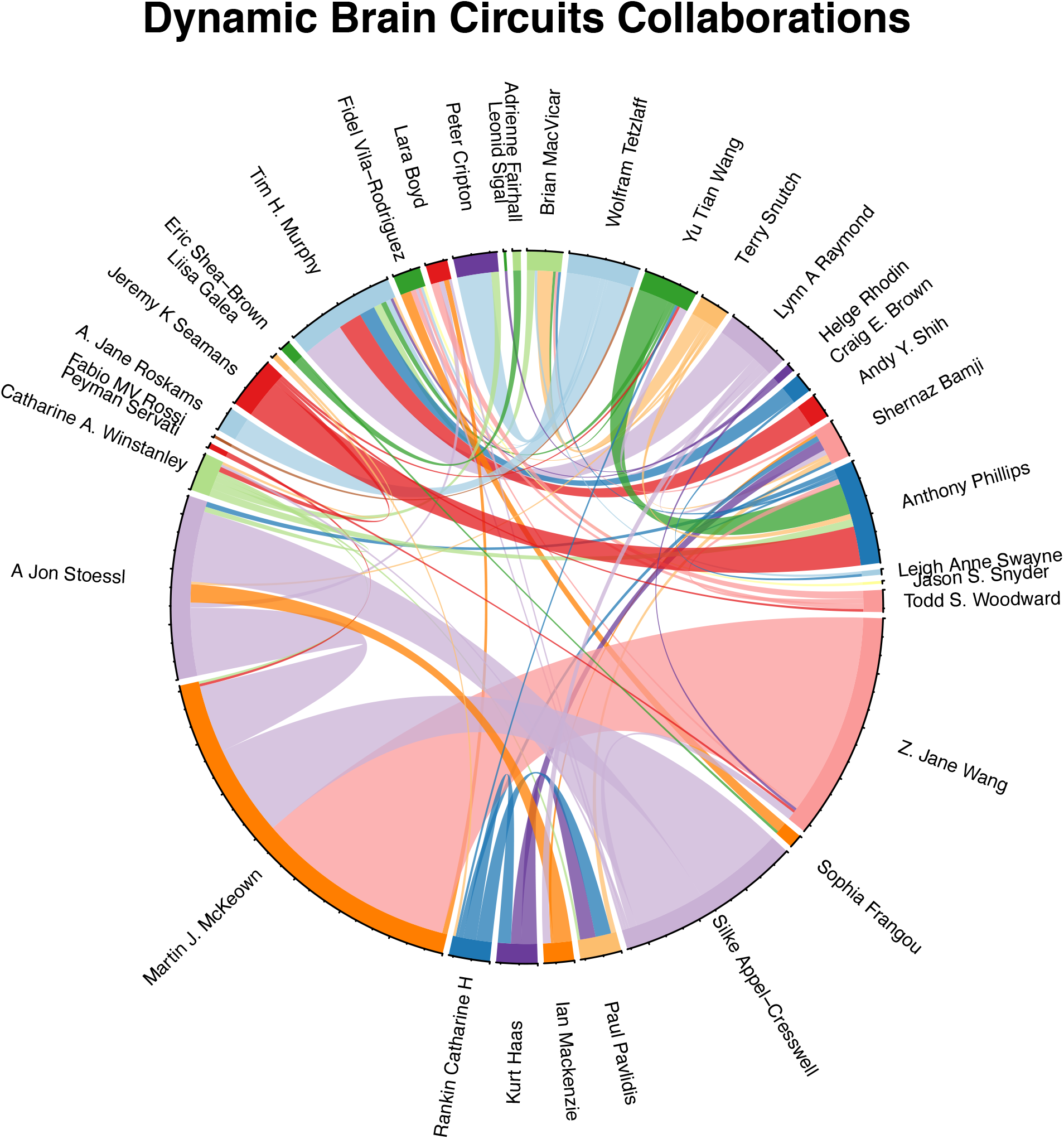
Collaboration chord diagram produced by the ScholarCollabs notebook. Links are weighted by the number of collaborations. The ScholarCollabs notebook creates this collaboration diagram based off of the Name and Coauthors columns (see Appendix C) of the author data table that is created as output by the ScholarScraper notebook.

**Fig. 5.**
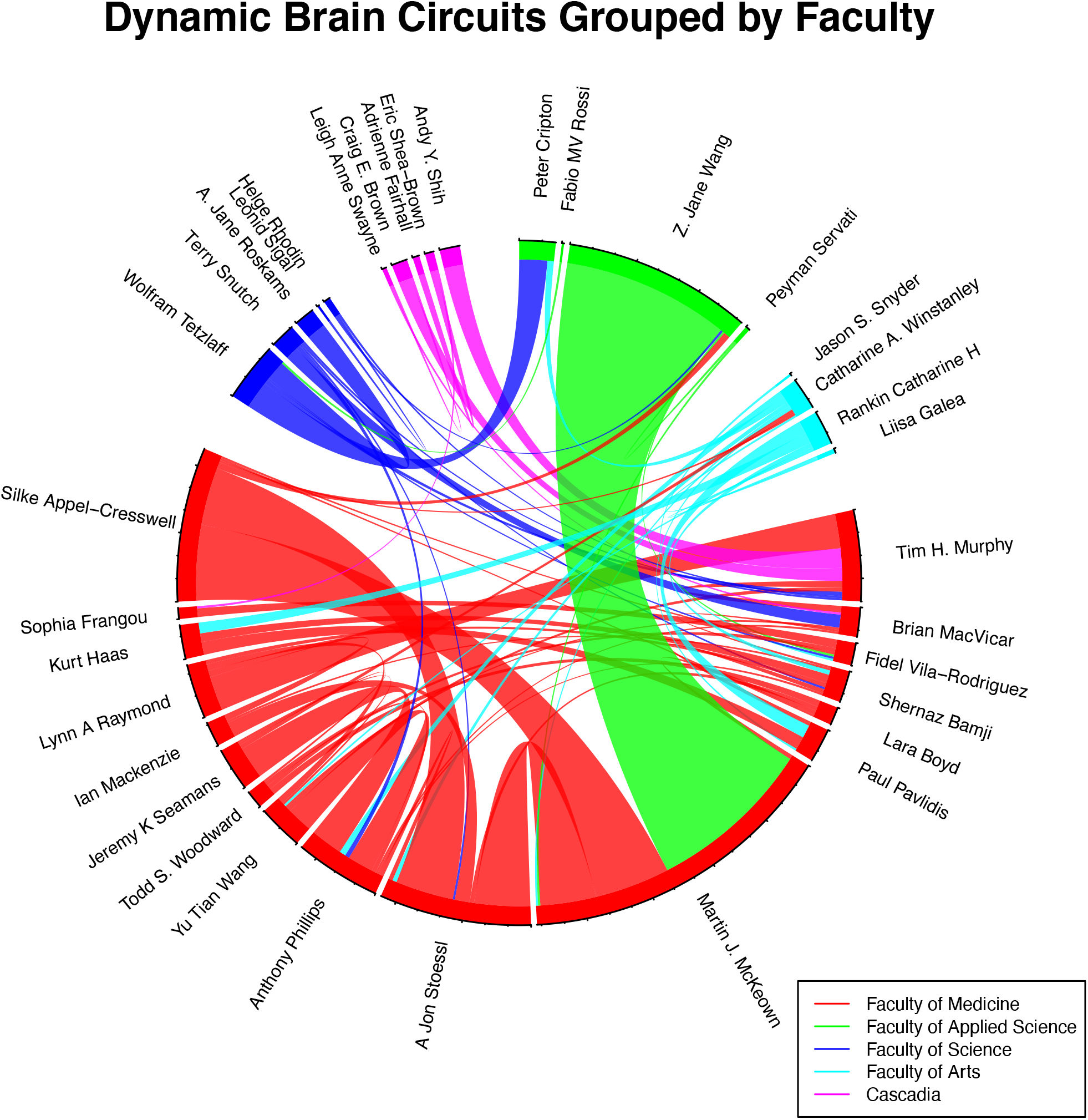
Grouped collaboration chord diagram created by the GroupedCollabs notebook based on faculty or institution. Links are weighted by the number of collaborations.

## 4. Impact

Bibliometric studies have investigated the strengths and limitations of a range of metrics as measures of research impact (Agarwal et al., 2016). For example, using metrics alone may favour some areas of research over others (Buchan et al., 2016). Other studies use bibliometric data to highlight the research topics in a field with the most citations, publication topic trends, and strength of collaboration networks between researchers (Buchan et al., 2016; Devos & Ménard, 2020; Yeung et al., 2017). Our tool helps to advance specific research streams by collecting bibliometric data, and a range of indicators, for a group of investigators while doing so in an automated, transparent manner.

In our experience, researchers who want to find bibliometrics data (for strategic planning purposes, evaluating researchers for promotion, or applying for grants, for example) currently do so by manually searching for each researcher and tracking data, or by using commercial packages such as SciVal. Manual methods are time consuming and error-prone. By addressing the need for automating the gathering of metrics, we wanted to ensure accessibility and reproducibility in the spirit of open science. SMS is open source, does not require coding experience, and comes with a detailed guide to setup and run the notebooks. To date, the SMS has been presented to and used by five Research Excellence Clusters at the University of British Columbia to assess their collaborative activities and productivity. The tool will continue to be improved while being broadly distributed to the academic community.

## 5. Conclusion

By allowing users to retrieve a comprehensive range of bibliometric data from a list of author names, SMS automates the process of retrieving information about the number of publications and key indicators for a group of authors. SMS is designed to be an open-source tool, accessible to those without coding backgrounds, and customizable to the specific needs of other projects and research groups.

## Acknowledgements

This work was supported by resources made available through the Dynamic Brain Circuits cluster and the NeuroImaging and NeuroComputation Centre at the UBC Djavad Mowafaghian Centre for Brain Health (RRID SCR_019086) and made use of the DataBinge forum. Timothy H Murphy (THM) was supported by Canadian Institutes of Health Research (CIHR) Foundation Grant FDN-143209. THM was also supported by the Brain Canada Neurophotonics Platform, a Heart and Stroke Foundation of Canada grant in aid, and a Leducq Foundation grant. We thank co-op students Carlos Doebeli, Ashli Forbes, and Keegan Flanagan for initial development work on this tool as well as the developers of the scholarly Python package. We thank the Pacific Institute for Mathematical Sciences for creating and maintaining the Syzygy.ca JupyterHub and the UBC Research Excellence clusters that helped evaluate and test early versions of SMS.

## Notes

### Competing Interest Statement

The authors have declared no competing interest.

