## Appendix A: Table 1 for "Scholar Metrics Scraper (SMS): automated retrieval of citation and author data"

**Table 1.** UBC Dynamic Brain Circuits Cluster author list. This is an example of the list required as input to the ScholarScraper notebook (as CSV). Note that the list contains a mix of author names and Scholar IDs which are both supported as inputs to the ScholarScraper notebook. Scholar IDs have been highlighted.

| |
|---|
| Tim H. Murphy |
| Annie Ciernia |
| Brian MacVicar |
| Fidel Vila-Rodriguez |
| Shernaz Bamji |
| Lara Boyd |
| Paul Pavlidis |
| Martin McKeown |
| A Jon Stoessl |
| Peter Cripton |
| Jason Snyder |
| Wolfram Tetzlaff |
| Anthony Phillips |
| Catharine Winstanley |
| Yu Tian Wang |
| Todd Woodward |
| Jeremy Seamans |
| mTkQFQ8AAAAJ |
| Ian Mackenzie |
| Lynn Raymond |

Kurt Haas

Mark Cembrowski

Fabio Rossi

Jane Roskams

Catharine Rankin

Michael Gordon

Leonid Sigal

W75uTm8AAAAJ

Peyman Servati

Liisa Galea

Sophia Frangou

Silke Cresswell

Helge Rhodin

Manu Madhav

Brian D. Fisher

Leigh Anne Swayne

Craig E. Brown

Adrienne Fairhall

Eric Shea-Brown

Emily Sylwestrak

Andy Shih
