## Appendix B: Table 2 for "Scholar Metrics Scraper (SMS): automated retrieval of citation and author data"

**Table 2.** Author data table. This is a portion of the table created by the ScholarScraper notebook as output (as CSV). Affiliations were flagged if they did not match one of: University of British Columbia, Simon Fraser University, University of Victoria, or University of Washington. The full version of this table will also contain the list of publication titles and coauthors.

| Name | Document Count | Cited by | h-index | i10-index | Affiliation | Warning |
|---|---|---|---|---|---|---|
| **Tim H. Murphy** | 226 | 18646 | 72 | 144 | University of British Columbia | |
| **Annie Vogel Ciernia** | 37 | 1456 | 17 | 22 | University of British Columbia, Vancouver | |
| **Brian MacVicar** | 204 | 18402 | 74 | 133 | Professor, University of British Columbia | |
| **Fidel Vila-Rodriguez** | 225 | 3575 | 27 | 62 | Assistant Professor; University of British Columbia | |
| **Shernaz Bamji** | 55 | 4634 | 30 | 37 | Professor, University of British Columbia | |
| **Lara Boyd** | 320 | 11385 | 58 | 143 | Professor, University of British Columbia | |
| **Paul Pavlidis** | 244 | 15463 | 67 | 136 | Professor of Psychiatry, University of British Columbia | |
| **Martin J. McKeown** | 346 | 14100 | 42 | 123 | Professor of Medicine (Neurology), University of British Columbia | |
| **A Jon Stoessl** | 452 | 26874 | 73 | 225 | University of British Columbia | |
| **Peter Cripton** | 287 | 6316 | 41 | 105 | Professor of Biomedical Engineering, University of British Columbia | |
| **Jason S. Snyder** | 34 | 5270 | 18 | 24 | Associate Professor, Department of Psychology, Djavad Mowafaghian Centre for Brain Health | |
| **Wolfram Tetzlaff** | 309 | 21418 | 74 | 175 | Professor and Director ICORD, University of British Columbia | |
| **Anthony Phillips** | 435 | 33992 | 100 | 284 | University of British Columbia | |
| **Catharine A. Winstanley** | 176 | 9608 | 45 | 86 | University of British Columbia | |
| **Yu Tian Wang** | 199 | 36636 | 74 | 140 | University of British Columbia | |
| **Todd S. Woodward** | 288 | 11706 | 61 | 147 | Professor, University of British Columbia | |
| **Jeremy K Seamans** | 104 | 14474 | 48 | 63 | University of British Columbia | |
| **Terry Snutch** | 455 | 28635 | 87 | 205 | University of British Columbia | |
| **Ian Mackenzie** | 470 | 49818 | 98 | 211 | Division of Neuropathology, Department of Pathology and Lab Medicine, University of British Columbia | |

| | | | | | | |
|---|---|---|---|---|---|---|
| **Lynn A Raymond** | 167 | 17472 | 69 | 118 | Professor of Psychiatry, University of British Columbia, Huntington Study Group, Predict-HD | |
| **Kurt Haas** | 71 | 3357 | 24 | 35 | Full Professor of Neuroscience, University of British Columbia | |
| **Mark S. Cembrowski** | 32 | 1429 | 16 | 17 | Assistant Professor, University of British Columbia | |
| **Fabio MV Rossi** | 125 | 14233 | 50 | 89 | Professor of Medical Genetics, University of British Columbia | |
| **A. Jane Roskams** | 69 | 5776 | 39 | 50 | University of British Columbia/Washington | |
| **Rankin Catharine H** | 207 | 6070 | 37 | 76 | University of British Columbia | |
| **Michael Gordon** | 35 | 3542 | 19 | 20 | University of British Columbia | |
| **Leonid Sigal** | 197 | 10468 | 48 | 103 | Associate Professor, University of British Columbia | |
| **Z. Jane Wang** | 405 | 9698 | 49 | 167 | Professor of Electrical and Computer Engineering Dept., University of British Columbia, Canada | |
| **Peyman Servati** | 265 | 7244 | 41 | 99 | Electrical and Computer Engineering, University of British Columbia | |
| **Liisa Galea** | 264 | 18917 | 74 | 156 | Professor, Department of Psychology, University of British Columbia | |
| **Sophia Frangou** | 606 | 22507 | 80 | 225 | Icahn School of Medicine at Mount SInai | Affiliation does not match! |
| **Silke Appel-Cresswell** | 96 | 3334 | 21 | 27 | Associate Professor, University of British Columbia | |
| **Helge Rhodin** | 55 | 2499 | 20 | 23 | Assistant Professor at UBC | |
| **Manu S Madhav** | 19 | 290 | 8 | 8 | Assistant Professor, University of British Columbia | |
| **Brian D. Fisher** | 176 | 3042 | 30 | 49 | Professor of Interactive Arts and Technology, Simon Fraser University | |
| **Leigh Anne Swayne** | 62 | 1625 | 23 | 35 | Associate Professor, University of Victoria | |
| **Craig E. Brown** | 32 | 2485 | 21 | 28 | University of Victoria, University of British Columbia | |
| **Adrienne Fairhall** | 109 | 4856 | 34 | 50 | Professor of Physiology and Biophysics, University of Washington | |
| **Eric Shea-Brown** | 189 | 6514 | 35 | 67 | Applied Mathematics, University of Washington | |
| **Emily Sylwestrak** | 11 | 1308 | 8 | 8 | Stanford University | Affiliation does not match! |
| **Andy Y. Shih** | 78 | 6100 | 33 | 46 | Seattle Children's Research Institute & University of Washington | |
