## Appendix C: Table 3 for "Scholar Metrics Scraper (SMS): automated retrieval of citation and author data"

**Table 3.** Coauthor data table. This is a portion of the table created by the ScholarScraper notebook as output (as CSV). These two columns are used as input for the ScholarCollab and GroupedCollabs notebooks to produce the collaboration diagrams (Fig. 4 and Fig. 5).

| Name | Coauthors |
|---|---|
| **Tim H. Murphy** | {'Andy Y. Shih': 9, 'Lynn A Raymond': 16, 'Craig E. Brown': 6, 'Snutch': 1, 'Yu Tian Wang': 2, 'Brian MacVicar': 3, 'Wolfram Tetzlaff': 2, 'Helge Rhodin': 1} |
| **Annie Vogel Ciernia** | |
| **Brian MacVicar** | {'Tim H. Murphy': 3, 'Snutch': 6, 'Anthony Phillips': 1, 'Shernaz Bamji': 1, 'Yu Tian Wang': 1, 'Leigh Anne Swayne': 1} |
| **Fidel Vila-Rodriguez** | {'Z. Jane Wang': 1, 'Sophia Frangou': 4, 'Silke Appel-Cresswell': 1, 'Jason S. Snyder': 1, 'Wolfram Tetzlaff': 1, 'Todd S. Woodward': 2} |
| **Shernaz Bamji** | {'Brian MacVicar': 1, 'Ian Mackenzie': 1, 'Lynn A Raymond': 2, 'Anthony Phillips': 1, 'Snutch': 1, 'Paul Pavlidis': 2, 'Kurt Haas': 4, 'Rankin Catharine H': 2} |
| **Lara Boyd** | {'Todd S. Woodward': 3, 'Martin J. McKeown': 2, 'A Jon Stoessl OR jon Stoessl': 2, 'Silke Appel-Cresswell': 1} |
| **Paul Pavlidis** | {'Shernaz Bamji': 2, 'Kurt Haas': 6, 'Rankin Catharine H': 6, 'Catharine A. Winstanley': 1} |
| **Martin J. McKeown** | {'Lara Boyd': 2, 'A Jon Stoessl OR jon Stoessl': 26, 'Silke Appel-Cresswell': 35, 'Z. Jane Wang': 77, 'Peyman Servati': 1, 'Catharine A. Winstanley': 1} |
| **A Jon Stoessl** | {'Lara Boyd': 2, 'Martin J. McKeown': 26, 'Silke Appel-Cresswell': 27, 'Catharine A. Winstanley': 2, 'Anthony Phillips': 2, 'Snutch': 1, 'Ian Mackenzie': 7} |
| **Peter Cripton** | {'Wolfram Tetzlaff': 13, 'Catharine A. Winstanley': 3} |
| **Jason S. Snyder** | {'Fidel Vila-Rodriguez': 1} |
| **Wolfram Tetzlaff** | {'Tim H. Murphy': 2, 'Fidel Vila-Rodriguez': 1, 'Peter Cripton': 13, 'Fabio MV Rossi': 1, 'A. Jane Roskams': 9} |
| **Anthony Phillips** | {'Brian MacVicar': 1, 'Shernaz Bamji': 1, 'A Jon Stoessl OR jon Stoessl': 2, 'Jeremy K Seamans': 15, 'Yu Tian Wang': 12, 'Todd S. Woodward': 2, 'Catharine A. Winstanley': 3, 'Snutch': 2} |
| **Catharine Winstanley** | {'Paul Pavlidis': 1, 'Martin J. McKeown': 1, 'A Jon Stoessl OR jon Stoessl': 2, 'Silke Appel-Cresswell': 1, 'Peter Cripton': 3, 'Anthony Phillips': 3, 'Jeremy K Seamans': 2, 'Liisa Galea': 1} |
| **Yu Tian Wang** | {'Tim H. Murphy': 2, 'Brian MacVicar': 1, 'Anthony Phillips': 12, 'Jeremy K Seamans': 1, 'Lynn A Raymond': 3, 'Rankin Catharine H': 1} |
| **Todd S. Woodward** | {'Fidel Vila-Rodriguez': 2, 'Lara Boyd': 3, 'Anthony Phillips': 2, 'Jeremy K Seamans': 1} |
| **Jeremy K Seamans** | {'Anthony Phillips': 15, 'Catharine A. Winstanley': 2, 'Yu Tian Wang': 1, 'Todd S. Woodward': 1} |
| **Snutch** | {'Tim H. Murphy': 1, 'Brian MacVicar': 6, 'Shernaz Bamji': 1, 'A Jon Stoessl OR jon Stoessl': 1, 'Anthony Phillips': 2} |
| **Ian Mackenzie** | {'Shernaz Bamji': 1, 'Lynn A Raymond': 3, 'A Jon Stoessl OR jon Stoessl': 7} |
| **Lynn A Raymond** | {'Tim H. Murphy': 16, 'Shernaz Bamji': 2, 'Ian Mackenzie': 3, 'Yu Tian Wang': 3} |
| **Kurt Haas** | {'Shernaz Bamji': 4, 'Paul Pavlidis': 6, 'Rankin Catharine H': 5} |

| | |
|---|---|
| **Mark S. Cembrowski** | |
| **Fabio MV Rossi** | {'Wolfram Tetzlaff': 1} |
| **A. Jane Roskams** | {'Wolfram Tetzlaff': 9} |
| **Rankin Catharine H** | {'Shernaz Bamji': 2, 'Paul Pavlidis': 6, 'Kurt Haas': 5, 'Yu Tian Wang': 1, 'Liisa Galea': 1} |
| **Michael Gordon** | |
| **Leonid Sigal** | {'Helge Rhodin': 1} |
| **Z. Jane Wang** | {'Fidel Vila-Rodriguez': 1, 'Martin J. McKeown': 77, 'Silke Appel-Cresswell': 3, 'Peyman Servati': 1, 'Helge Rhodin': 1} |
| **Peyman Servati** | {'Martin J. McKeown': 1, 'Z. Jane Wang': 1} |
| **Liisa Galea** | {'Catharine A. Winstanley': 1, 'Rankin Catharine H': 1} |
| **Sophia Frangou** | {'Fidel Vila-Rodriguez': 4, 'Eric Shea-Brown': 1} |
| **Silke Appel-Cresswell** | {'Fidel Vila-Rodriguez': 1, 'Lara Boyd': 1, 'Martin J. McKeown': 35, 'A Jon Stoessl OR jon Stoessl': 27, 'Z. Jane Wang': 3, 'Catharine A. Winstanley': 1} |
| **Helge Rhodin** | {'Tim H. Murphy': 1, 'Leonid Sigal': 1, 'Z. Jane Wang': 1} |
| **Manu S Madhav** | |
| **Brian D. Fisher** | |
| **Leigh Anne Swayne** | {'Brian MacVicar': 1, 'Craig E. Brown': 1} |
| **Craig E. Brown** | {'Tim H. Murphy': 6, 'Leigh Anne Swayne': 1} |
| **Adrienne Fairhall** | {'Eric Shea-Brown': 3} |
| **Eric Shea-Brown** | {'Sophia Frangou': 1, 'Adrienne Fairhall': 3} |
| **Emily Sylwestrak** | |
| **Andy Y. Shih** | {'Tim H. Murphy': 9} |
