## Appendix D: Source Code 1 for "Scholar Metrics Scraper (SMS): automated retrieval of citation and author data"

**Source Code 1:** ScholarScraper notebook

THE UNIVERSITY OF BRITISH COLUMBIA

Dynamic Brain Circuits in Health & Disease

Research Excellence Cluster

### 1 Scholar Metrics Scraper: ScholarScraper notebook

##### Introduction

This notebook automates the process of retrieving bibliometric data from Google Scholar for a list of authors. It utilizes scholarly, a Python module that allows users to retrieve bibliometrics from Google Scholar. It also creates a citations/year bar chart and a collaborations heatmap. This notebook should be run before the ScholarCollabs and GroupedCollabs notebooks.

This project currently works with scholarly 1.4.5

##### Installation and Setup

1. Set-up Jupyter. If your institution has access, you can use Syzygy to run in the Cloud, or install on your computer following these instructions.
2. Clone the project.
    - Open Terminal (in Syzygy, click the "+" button to open a new launcher and click "Terminal"
    - Type "git clone https://github.com/ubcbraincircuits/scholar_metrics_scraper" and press enter
    - The project should now be cloned in your directory.
    - Alternatively, you can download the project as a ZIP file from https://github.com/ubcbraincircuits/scholar_metrics_scraper(click Code, then Download ZIP)
3. Install scholarly
    - In the terminal (from above) type "pip install scholarly" or "pip install –user scholarly" and press enter

4. Obtain a CSV file with the list of author names in a single column with no column header. Ideally, all author names should match their names in Google Scholar. Upload this file to Syzygy or move it to the project folder on your computer. This file should be in the same directory as this notebook file (ScholarScraper.ipynb).
5. Modify the names of the input/output files below (in step 2). The input file name must match the CSV file name.
6. Modify the "affiliations" variable as a list of institution names which the researchers are affiliated with. Include both abbreviated and long form.

7. Run all cells (click shift+enter to run a cell or the play button above).
8. Open the ouput CSV file in the same directory as this notebook file. Check the last column of this file for warnings. If needed, modify the author names in the input CSV file if the wrong author profile was scraped, or no profile was found, and re-run.

**Steps**

1. Install and load libraries and packages.

[ ]:
```
#If you receive an error running this cell for the first time, try running it␣
↪agian.

from scholarly import scholarly
import csv
import warnings
import pandas as pd
import numpy as np
import matplotlib
from matplotlib import pyplot as plt
from matplotlib import cm as CM
```

2. Modify the names of the input and output files. The name of the input file should match the name of the author list CSV file. If you followed the setup instructions, the CSV file should now be in the same directory as this notebook file. The output file does not have to exist yet (it will be created).

[ ]:
```
# !!! Modify this to match the name of your author list CSV file.
author_list_csv = 'authorlist.csv'

output_data_csv = 'ss_output_data.csv'
```

3. Load in the author names from the CSV file.

[ ]:
```
author_names= []
with open(author_list_csv, encoding ="utf-8-sig") as csv_file:
    csv_reader = csv.reader(csv_file, delimiter =',')
    for row in csv_reader:
        if (len(row) == 1):
            author_names.append(row[0])
```

4. Modify the affiliations list with institutions which the researchers are affiliated with.

```
[ ]: affiliations = ['University of British Columbia', 'UBC', 'Djavad Mowafaghian',␣
     ↪'Simon Fraser University',
                     'University of Victoria', 'University of Washington']
```

5. Scrape data for each author. This will take several minutes.

```
[ ]: # This will contain all the data for each author which will be exported as a␣
     ↪table. It will be a list of dictionaries.
     rows = []
     # This will contain a list of dictionaries for each author. The dictionaries␣
     ↪will be made up of years as keys and citation numbers as vals
     cites_per_year = []
     # This dictionary will contain publication titles as keys and author names as␣
     ↪values
     pub_authors = {}


     for athr in author_names:
         pubs = []
         try :
             if ' ' in athr:
                 search_query = scholarly.search_author(athr)
                 author = next(search_query)
             else:
                 author = scholarly.search_author_id(athr)
         except (RuntimeError,TypeError,StopIteration):
             row = {'Name': athr, 'Warning': 'no information found'}
         else:
             data_dict = scholarly.fill(author, sections=['basics', 'indices',␣
     ↪'publications', 'counts'])

             # Get publications titles
             for pub in data_dict['publications']:
                 pubs.append(pub['bib']['title'])
                 # Add to dictionary with title as key and author as value
                 pub_authors.setdefault(pub['bib']['title'],[]).
     ↪append(data_dict['name'])

             # Get citations per year and put in dictionary
             cites_per_year_dict = data_dict['cites_per_year']
             # Add name to dictionary
             cites_per_year_dict['name'] = data_dict['name']
             cites_per_year.append(cites_per_year_dict)

             # Create row (dictionary) for output data table
```

```
        row = {'Name': data_dict['name'], 'Scholar ID': data_dict['scholar_id'],
               'Cited by': data_dict['citedby'], 'Cited by 5 years':␣
↪data_dict['citedby5y'],
               'h-index': data_dict['hindex'], 'h-index 5 years':␣
↪data_dict['hindex5y'],
               'i10-index': data_dict['i10index'],'i10-index 5 years':␣
↪data_dict['i10index5y'],
               'Publications': pubs, 'Document Count': len(pubs), 'Affiliation':
↪ data_dict['affiliation']}

        # Create list of authors who do not have the specified affiliation
        if not any(a in data_dict['affiliation'] for a in affiliations):
            row['Warning'] = "Affiliation does not match!"

    finally:
        rows.append(row)
```

6. Add coauthors to the rows.

```
[ ]: # Create dictionary with author names as keys and dictionary (coauthor name as␣
↪key, number of collaborations as value)
#   as value
collabs_dict={}
for key in pub_authors:
    for author in pub_authors[key]:
        for coauthor in pub_authors[key]:
            if coauthor is not author:
                if author not in collabs_dict.keys() or coauthor not in␣
↪collabs_dict[author].keys():
                    collabs_dict.setdefault(author,{})[coauthor]=1
                else:
                    collabs_dict[author][coauthor]+=1


# Write to rows dataframe
for row in rows:
    if row['Name'] in collabs_dict.keys():
        row['Coauthors'] = collabs_dict[row['Name']]
```

7. Write rows to output CSV file

[ ]:
```
# Specify the order of the data. These keys must match the names of the keys in␣
↪the rows dictionary.
keys = ['Name', 'Scholar ID', 'Document Count', 'Cited by', 'Cited by 5 years',␣
↪'h-index', 'h-index 5 years',  'i10-index', 'i10-index 5 years',␣
↪'Publications', 'Coauthors', 'Affiliation', 'Warning']

# This creates/opens the file with filename with the intention to write to the␣
↪csv_file
# The encoding allows the characters to be properly written to the csv_file
with open(output_data_csv, mode='w', encoding ="utf-8") as csv_file:
    csv_writer = csv.writer(csv_file, delimiter=',', quotechar='"', quoting=csv.
↪QUOTE_MINIMAL)
    dict_writer = csv.DictWriter(csv_file, keys, delimiter=',', quotechar='"',␣
↪quoting=csv.QUOTE_MINIMAL)
    dict_writer.writeheader()
    dict_writer.writerows(rows)
```

8. Create barplot of citations per year. Modify the years and output PDF filename.

[ ]:
```
# Create citations per year dataframe
cites_df = pd.DataFrame(cites_per_year)

# Add a totals column
cites_df.loc['Total']= cites_df.sum()

# Select years to plot
cites_df_selected = cites_df[[2016,2017,2018,2019,2020,2021]]

# Select the last row (totals)
cites_df_total = cites_df_selected.iloc[-1:]

# Create barplot
years = list(cites_df_total.columns)
cites = cites_df_total.values.tolist()[0]
plt.bar(years, cites )
plt.ylabel('Total Citations')

# !!! Modify this - name the output PDF file
plt.savefig("citations.pdf")
```

9. Create collaboration heatmap

[ ]:
```
authors = list(collabs_dict.keys())
author_count=len(authors)
# Initialize array with zeros.
collabs_array = np.zeros((author_count, author_count), dtype=int)
```

```python
# Populate the array with the number of collaborations between authors
for i, athr in enumerate(authors):
    for j, name in enumerate(authors):
        if name in collabs_dict[athr].keys():
            collabs_array[i][j] = collabs_dict[athr][name]

# Only display the lower triangle of the matrix
mask =  np.tri(collabs_array.shape[0], k=-1)
collabs_array = np.ma.array(collabs_array, mask=mask).T

# Set up the colourmap
cmap = CM.get_cmap('nipy_spectral')
cmap = cmap.reversed()

### Create heatmap ###

fig = plt.figure(figsize = (author_count,author_count))
ax = fig.add_subplot(111)
im = ax.imshow(collabs_array, cmap=cmap, interpolation='nearest',
               #norm=matplotlib.colors.LogNorm()            # Uncomment this if␣
 ↪you want a logarithmic colormap
              )

# Show the ticks
ax.set_xticks(np.arange(author_count))
ax.set_yticks(np.arange(author_count))

# Label the ticks with author names - you can modify the authorname font size␣
 ↪here
ax.set_xticklabels(authors,fontsize=25)
ax.set_yticklabels(authors, fontsize=25)

# Rotate and align tick labels.
plt.setp(ax.get_xticklabels(), rotation=45, ha="right",
         rotation_mode="anchor")

# Create numeric annotations.
for i in range(author_count):
    for j in range(author_count):
        text = ax.text(j, i, collabs_array[i, j],
                       ha="center", va="center", color="w", fontsize=25)
# Fit plot within figure
fig.tight_layout()


# Add title - you can change the title and title font size here
ax.set_title("Publication Collaborations", fontsize = 50)
```

```python
# Add the colorbar and add ticks - change colorbar label fontsize here
cbar = fig.colorbar(im)
cbar.ax.tick_params(labelsize=30)

# Save the figure as PDF - you can modify the filename here.
plt.savefig("collabs_heatmap.pdf")

plt.show()
```
