## Appendix E: Source Code 2 for "Scholar Metrics Scraper (SMS): automated retrieval of citation and author data"

**Source Code 2:** ScholarCollabs notebook

THE UNIVERSITY OF BRITISH COLUMBIA

Dynamic Brain Circuits in Health & Disease
Research Excellence Cluster

### 1 Scholar Metrics Scraper: ScholarCollabs notebook

##### Introduction

This R notebook creates a chord diagram to visualize publication collaborations between coauthors. It creates links between authors and their coauthors from the output CSV file created by the ScholarScraper.ipynb notebook. This notebook should be run after you run the ScholarScraper notebook. This notebook does not group the authors into sub-groups. For this purpose, use GroupedCollabs.ipynb.

**Installation and Setup** 1. At this point we will assume you have this project loaded in Jupyter and have successfully run the ScholarScraper notebook, which has created an output CSV file with author data. If this is not the case, go to the ScholarScraper.ipynb file and follow the instructions to setup and run.

2. Ensure that the CSV file created by ScholarScraper is in the same directory as this notebook. Make sure it has a column labeled 'Name' and a column labeled 'Coauthors'.

##### Steps

1. Install and load libraries

[ ]:
```
library(tidyverse)
library(viridis)
devtools::install_github("thomasp85/patchwork")
devtools::install_github("jokergoo/circlize")
install.packages("RColorBrewer")
library(patchwork)
```

```
library(circlize)
library(RColorBrewer)
```

2. Define the name of the data file (CSV created by Scholar Scraper)

[ ]:
```
# !!! Modify this to match the name of the CSV file that was created as output␣
↪form the ScholarScraper notebook
ss_output_file = "ss_output_data.csv"
```

3. Define the title, colors, and whether you want to create weighted or non-weighted diagram.

   View color palettes here

[ ]:
```
# !!! Modify the diagram title
title = "Dynamic Brain Circuits Collaborations"

# !!! Modify the colour palette.
c_pallete <- brewer.pal(12,"Paired")

# !!! Modify this - Set to TRUE if you want a weighted diagram or FALSE if you␣
↪want a non-weighted diagram.
weighted = TRUE
```

4. Load in collaboration data.

[ ]:
```
df_collab = read.csv(ss_output_file)
```

5. Tidy the dataframe

[ ]:
```
# creating a subset of our survey data that extracts the useful columns.
df_collab = subset(df_collab, select = c(Name, Coauthors))

# get rid of rows containing NAs
df_collab=df_collab[rowSums(is.na(df_collab)) != ncol(df_collab), ]
```

6. Set up the links

[ ]:
```
origin = c()
destination = c()
count = c()

for (i in 1:nrow(df_collab)) {
  x = df_collab$Name[i]
  for (n in 1:nrow(df_collab)) {
    if(is.na(df_collab$Coauthors[n]) == FALSE) {
      if(str_detect(df_collab$Coauthors[n], x) == TRUE) {
        origin = append(origin, df_collab$Name[n])
        destination = append(destination, x)
```

```
#       extracts digits that come after author name in␣
↪df_collab$Collaborators[n]
        count = append(count, strtoi(
            str_extract(df_collab$Coauthors[n],paste("(?<=", df_collab$Name[i],␣
↪"\': )\\d+", sep=""))))
      }
    }
  }
}

edge_l = data.frame(origin, destination, count)
# cleaning up the edge list by removing duplicates
edge_l = unique(edge_l)
edge_l$temp = apply(edge_l, 1, function(x) paste(sort(x), collapse=""))
edge_l = edge_l[!duplicated(edge_l$temp), 1:3]

 # create an adjacency list.
adjacencyData = data.frame(with(edge_l, table(origin, destination)))

# set the links to the edge list for a weighted diagram or adjacency list for␣
↪non-weighted
if (weighted == TRUE){
    links = edge_l
} else {
    links = adjacencyData
}
```

7. Assign a color to each investigator.

[ ]:
```
color =  c()

for (i in 1:nrow(df_collab)) {
    j=i%%12
    if(j==0){
        j= 12
        }
      color = append(color,c_pallete[j])
}

# adding the color column to the dataframe.
df_collab$color = color

# creating colour_ind structure
color_ind = structure(df_collab$color, names = df_collab$Name)
```

8. Create the chord diagram. Modify the name of the output PDF file. You can make additional optional modifications as well (read the comments below).

[ ]:

```
# Modify this!!!
pdf("chord_diagram.pdf")

# set up the parameters
circos.clear()
circos.par(start.degree = 90,gap.degree = 1,
           track.margin = c(-0.1, 0.1),
           points.overflow.warning = FALSE, canvas.xlim = c(-1.3,1.3),
           canvas.ylim = c(-1.3,1.3))
par(mar = c(0,0,2,0),xpd = TRUE, cex.main = 1.5)

# create the chord diagram
chordDiagram(links,
              grid.col = color_ind,
             transparency = 0.25,
             diffHeight  = -0.04,
             annotationTrack = "grid",
             annotationTrackHeight = c(0.05, 0.1),
             link.sort = TRUE,
             link.largest.ontop = FALSE,
              self.link = 1,
              small.gap = 1,
              big.gap = 1
)



# Add the text and the axis surrounding the diagram.
circos.trackPlotRegion(
  track.index = 1,
  bg.border = NA,
  panel.fun = function(x, y) {

    xlim = get.cell.meta.data("xlim")
    sector.index = get.cell.meta.data("sector.index")

    # Add names to the sector.
    # You can modify the font size of the names by changing cex and the␣
 ↪distance between the names
    #    and the circle by changing y.
    circos.text(
      x = mean(xlim),
      y = 6,
      labels = sector.index,
      facing = "clockwise",
      niceFacing = TRUE,
      cex = 0.7,
    )
```

```
     #Add graduation on axis
    circos.axis(
      h = "top",
      labels.cex = 0.001,
      minor.ticks = 2,
      major.tick.length = 0.1,
      labels.niceFacing = FALSE)

  }
)



# Add the title (user can modify the title in step 3)
title(title,outer=FALSE)


dev.off()
```
