## Appendix F: Source Code 3 for "Scholar Metrics Scraper (SMS): automated retrieval of citation and author data"

**Source Code 3:** GroupedCollabs notebook

THE UNIVERSITY OF BRITISH COLUMBIA

Dynamic Brain Circuits in Health & Disease
Research Excellence Cluster

### 1 Scholar Metrics Scraper: GroupedCollabs notebook

##### Introduction

This R notebook creates a **grouped** chord diagram to visualize publication collaborations between coauthors. It creates links between authors and their coauthors from the output CSV file created by the ScholarScraper.ipynb notebook. It also takes another CSV file as input to specify each author's group. This notebook should be run after the ScholarScraper notebook. If you do not want a grouped diagram, use ScholarCollabs.ipynb.

**Installation and Setup** 1. At this point we will assume you have this project loaded in Jupyter and have successfully run the ScholarScraper notebook, which has created an output CSV file with author data. If this is not the case, go to the ScholarScraper.ipynb file and follow the instructions to setup and run.

2. Ensure that the CSV file created by ScholarScraper is in the same directory as this notebook. Make sure it has a column labeled 'Name' and a column labeled 'Coauthors'.

3. Create a CSV file for the groupings. This should contain the author names in the first column, the group names as column names, and the author names under their respective groups (see the example below). The names in this file should match the names in the CSV file created by ScholarScraper. Upload this CSV to the project directory (the same directory as this notebook

file).

| | Group 1 | Group 2 | Group 3 |
|---|---|---|---|
| **Author 1** | Author 1 | | |
| **Author 2** | | Author 2 | |
| **Author 3** | | | Author 3 |
| **Author 4** | | Author 4 | |

**Steps**

1. Install and load libraries

[ ]:
```
library(tidyverse)
library(viridis)
devtools::install_github("thomasp85/patchwork")
devtools::install_github("jokergoo/circlize")
install.packages("RColorBrewer")
library(patchwork)
library(circlize)
library(RColorBrewer)
```

2. Define the name of the data file (CSV created by Scholar Scraper), investigator names CSV file, and group CSV file

[ ]:
```
# !!! Modify this to match the name of the CSV file that was created as output␣
↪form the ScholarScraper notebook
ss_output_file = "ss_output_data.csv"
# !!! Modify this to match the name of your groupings CSV file. See␣
↪instructions above.
group_file = "dbc_faculty_groups.csv"
```

3. Define the title, colors, and whether you want to create weighted or non-weighted diagram.

   View color options here.

[ ]:
```
# !!! Modify the diagram title
title = "Dynamic Brain Circuits Grouped by Faculty"

# !!! Modify the colour palette. Make sure there are the same number of colours␣
↪as groups.
c_pallete = c("red","green","blue","cyan","magenta")

# !!! Modify the group names. These groups will be paired with the colours in␣
↪the c_palette, in the same order.
group_names = c("Faculty of Medicine",
                               "Faculty of Applied Science",
                               "Faculty of Science",
                               "Faculty of Arts",
                               "Cascadia")
```

```
# !!! Modify this - Set to TRUE if you want a weighted diagram or FALSE if you
↪want a non-weighted diagram.
weighted = TRUE
```

4. Load in collaboration data

```
[ ]: df_collab = read.csv(ss_output_file)
```

5. Tidy the df_collab dataframe

```
[ ]: # creating a subset of our survey data that extracts the useful columns.
df_collab = subset(df_collab, select = c(Name, Coauthors))

# get rid of rows containing NAs
df_collab=df_collab[rowSums(is.na(df_collab)) != ncol(df_collab), ]
```

6. Set up the links

```
[ ]: ##### create an edge list using for loop. ####
origin = c()
destination = c()
count = c()

for (i in 1:nrow(df_collab)) {
  x = df_collab$Name[i]
  for (n in 1:nrow(df_collab)) {
    if(is.na(df_collab$Coauthors[n]) == FALSE) {
      if(str_detect(df_collab$Coauthors[n], x) == TRUE) {
        origin = append(origin, df_collab$Name[n])
        destination = append(destination, x)
#       extracts digits that come after author name in df_collab$Coauthors[n]
        count = append(count, strtoi(
            str_extract(df_collab$Coauthors[n],paste("(?<=", df_collab$Name[i],
↪"\': )\\d+", sep=""))))
      }
    }
  }
}

edge_l = data.frame(origin, destination, count)
# cleaning up the edge list by removing duplicates
edge_l = unique(edge_l)
edge_l$temp = apply(edge_l, 1, function(x) paste(sort(x), collapse=""))
edge_l = edge_l[!duplicated(edge_l$temp), 1:3]

# create an adjacency list.
adjacencyData = data.frame(with(edge_l, table(origin, destination)))
```

```
# set the links to the edge list for a weighted diagram or adjacency list for␣
↪non-weighted
if (weighted == TRUE){
    links = edge_l
} else {
    links = adjacencyData
}
```

7. Set up the groupings

[ ]:
```
# loading in the group names
df_group = read.csv(group_file)



# Setting up group names
Primary_c = colnames(df_group)[-1]

# Removing unnecessary first column
df_group <- df_group[,-1]

# Creating a group column by pivoting
df_group = df_group %>%
  pivot_longer(Primary_c,
               names_to = "group",
               values_to = "name")

# Removing NAs
df_group = na.omit(df_group)



# Assigning a colour to each group.
color =  c()

for (i in 1:nrow(df_group)) {
  for (j in 1:length(Primary_c)) {
    if(df_group$group[i] == Primary_c[j]) {
      color = append(color,c_pallete[j])
    }
  }
}

# adding the color column to the dataframe.
df_group$color = color

# creating the groupings
group_ind = structure(df_group$group, names = df_group$name)
```

```
# creating colors for the groupings
color_ind = structure(df_group$color, names = df_group$name)
```

8. Create the chord diagram. Modify the name of the output PDF file. You can make additional optional modifications as well (read the comments below).

[ ]:
```
# !!! Modify this
pdf("grouped_chord_diagram.pdf")

# set up the parameters
circos.clear()
circos.par(start.degree = 90,gap.degree = 1,
           track.margin = c(-0.1, 0.1),
           points.overflow.warning = FALSE, canvas.xlim = c(-1.3,1.3),
           canvas.ylim = c(-1.3,1.3))
par(mar = c(0,0,2,0),xpd = TRUE, cex.main = 1.5)

# create the chord diagram
chordDiagram(links, group = group_ind,
              grid.col = color_ind,
             transparency = 0.25,
             diffHeight  = -0.04,
             annotationTrack = "grid",
             annotationTrackHeight = c(0.05, 0.1),
             link.sort = TRUE,
             link.largest.ontop = FALSE,
              self.link = 1,

)



# Add the text and the axis surrounding the diagram.
circos.trackPlotRegion(
  track.index = 1,
  bg.border = NA,
  panel.fun = function(x, y) {

    xlim = get.cell.meta.data("xlim")
    sector.index = get.cell.meta.data("sector.index")

    # Add names to the sector.
    # You can modify the font size of the names by changing cex and the␣
↪distance between the names
    #    and the circle by changing y.
    circos.text(
      x = mean(xlim),
```

```
      y = 6,
      labels = sector.index,
      facing = "clockwise",
      niceFacing = TRUE,
      cex = 0.7,
    )

     #Add graduation on axis
    circos.axis(
      h = "top",
      labels.cex = 0.001,
      minor.ticks = 2,
      major.tick.length = 0.1,
      labels.niceFacing = FALSE)

  }
)
# Add a legend
# You can modify the position (e.g. change "bottomright" to "topleft"), and the␣
↪font (cex)
legend("bottomright", legend=group_names,
       col=c_pallete, lty=1, cex=0.7)


# Add the title (user can modify the title in step 3)
title(title,outer=FALSE)


dev.off()
```
